# *VIVIDHA*: *V*ar*i*ant Prediction and *V*isualization *I*nterface for *D*ynamic *H*igh-throughput *A*nalysis

**DOI:** 10.1101/2025.01.29.635418

**Authors:** Tarang Barasiya, Neeraj Bharti, Renu Gadhari, Avinash Bayaskar, Sankalp Jain, Amit Saxena, Sunitha Manjari Kasibhatla, Rajendra Joshi

## Abstract

Large scale genome sequencing projects have produced huge datasets that pose challenges of high processing times especially for variant calling, a significant downstream analysis step. Efficient utilization of computational resources for accurate variant prediction in a timely manner is possible using Hadoop MapReduce framework. We have developed VIVIDHA (***V***ar***i***ant Prediction and *V*isualization *I*nterface for *D*ynamic *H*igh-throughput Analysis), a high throughput methodology for prediction of variants based on splitting the alignment file using overlapping regions using Hadoop MapReduce framework. The size of overlap region is user-defined. Three variant callers viz. GATK, VarScan2 and BCFTools have been included to predict variants using a consensus approach. Speed-up observed provides the rationale of better performance as number of compute cores and file size are increased. VIVIDHA is available in both GUI as well as command-line modes and can be downloaded from URL: https://github.com/bioinformatics-cdac/VividhaInstaller

## 1. Introduction

With the advent of high throughput sequencing technologies there has been an explosive growth of sequencing data, which mandates availability of faster and more efficient data analysis tools [Schmidt and Hildebrandt 2017]. Variant calling is a key step in the next generation sequencing (NGS) data analysis workflow. The quality of call sets directly affects downstream analysis such as detection of disease-causing gene(s) [Krusche *et al*. 2019]. Joint variant calling is highly relevant for genotype-phenotype associations using resources like gnomAD [Karczewski *et al*. 2020] for precision public-health [Khoury *et al*. 2016]. With the increasing size of datasets, there is a dire need to develop scalable and cost-effective methodologies for variant calling. Existing solutions are based either on Hadoop MapReduce [Decap *et al*. 2016; Mohammed *et al*. 2014] or Spark [Xiao *et al*. 2019; Petrillo *et al*. 2021] frameworks including shell scripts [Huang *et al*. 2016; Kelly *et al*. 2015] and scalable persistent databases [Lin *et al*. 2018]. Majority of these solutions are associated with a steep learning curve for life sciences researchers as they need to be conversant with command-line usage of tools in linux environment. The objective of the present work is to develop an graphical user interface based computational platform that shields the users from complexities associated with using high performance compute clusters for variant calling of multiple samples at a single instance using the strategy of splitting the input alignment files with overlapping boundaries.

## 2. Methodology

VIVIDHA is based on distribution of alignment files (BAM) obtained after reference mapping into several overlapping chunks to enable fast computation without compromising on the quality of the called variants. Pre-processed alignment files (sorted BAM) are split into overlapping chunks using samtools v 1.8. Each of the partitioned dataset is then subjected to variant prediction using the MapReduce methodology. Variant prediction can then be carried out using three tools viz., GATK v 4.0[McKenna *et al*. 2010; DePristo *et al*. 2011], VarScan2 v 2.4.3[Koboldt *et al*. 2012] and BCFtools v 1.9 [Narasimhan *et al*. 2016] either in an independent manner or simultaneously followed by a consensus approach. Best results in terms of specificity for SNP calculation are achieved when strict consensus of all calls is applied (Figure 1).

**Figure 1:**
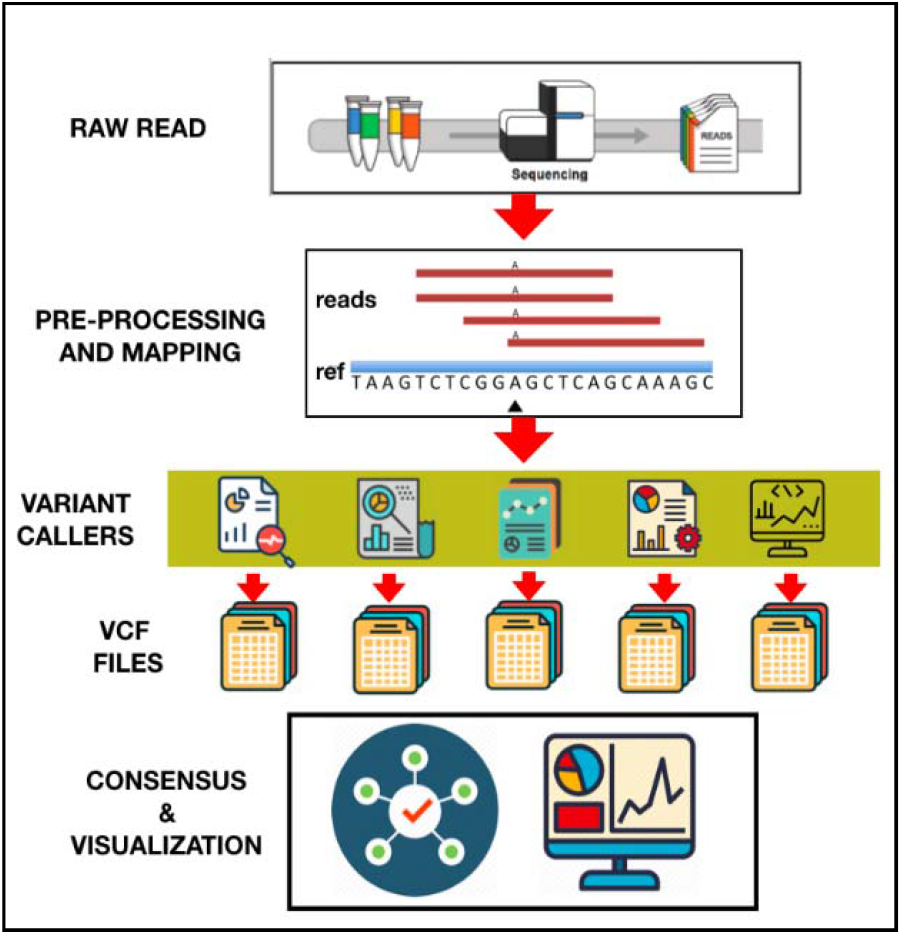
Flow-chart of VIVIDHA

VIVIDHA is based on Hadoop which is an Open Source Big data Solution framework for both distributed storage and distributed computing using clusters of commodity hardware. MapReduce programming model has been used for development of VIVIDHA application that can process big data in parallel on multiple nodes.

### 2.1 Map Reduce

MapReduce is a software framework and programming model used for processing huge amounts of data. MapReduce programs work in two phases, namely, Map and Reduce. Map tasks deal with splitting and mapping of data while Reduce tasks shuffle and reduce the data. The data goes through the following phases of MapReduce in Big Data (Figure 2):

**Figure 2:**
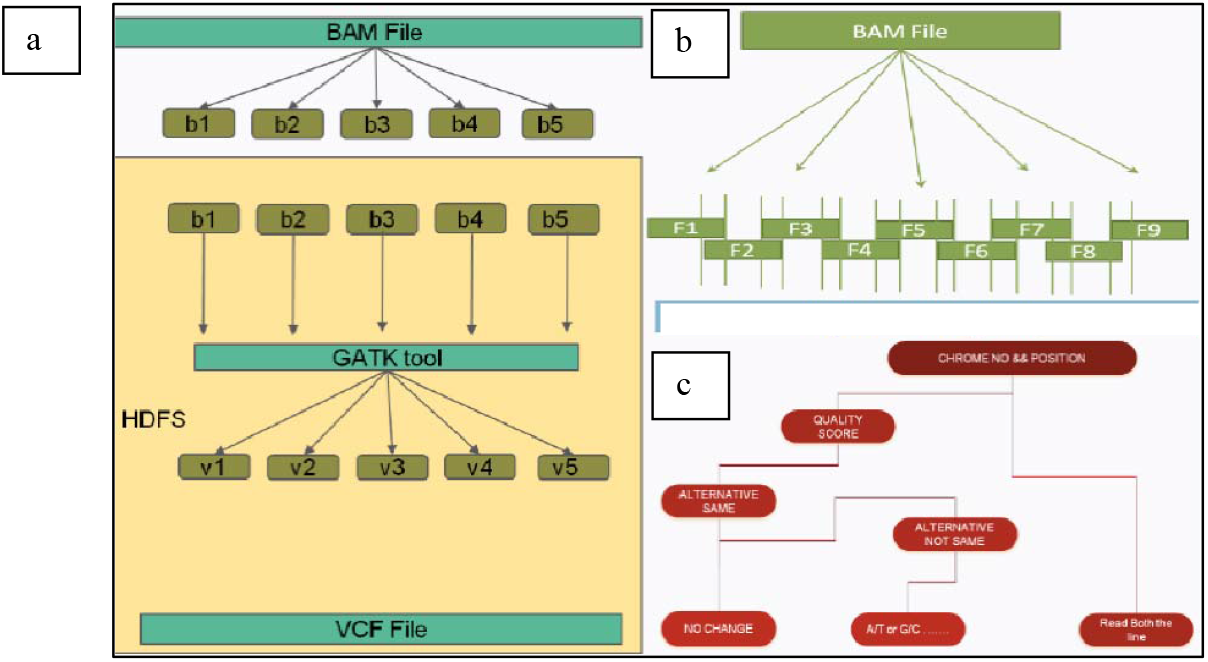
a) Flowchart for generating overlapping alignment files b) File splitting strategy c) Logic of reducer

#### 2.1.1 Input Splits

An input to a MapReduce in Big Data job is divided into fixed-size pieces called input splits which is a chunk of the input that is consumed by a single map.

#### 2.1.2 Mapping

This is the very first phase in the execution of map-reduce program. In this phase, data in each split is passed to a mapping function to produce output values. In the current implementation, a job of mapping phase is to count a number of occurrences of each word from input splits and prepare a list in the form of <word, frequency>

#### 2.1.3 Shuffling

This phase consumes the output of the Mapping phase. Its task is to consolidate the relevant records from Mapping phase output. In the current implementation, the same words are clubbed together along with their respective frequency.

#### 2.1.4 Reducing

In this phase, output values from the Shuffling phase are aggregated and a single output value is returned thereby summarizing the complete dataset.

#### 2.1.5 File Splitting

The alignment file (BAM) has been used as an Input file which is further divided into overlapping chunks using BAMTools. The BAM file is divided into different splits and different overlap areas. The first division of the BAM files is done chromosome-wise and after that division is into various splits with overlap.

### 2.2 Hadoop Architecture

Hadoop is a scalable, open source, fault-tolerant Virtual Grid operating system architecture for data storage and processing. It runs on commodity hardware and uses HDFS which is fault-tolerant high-bandwidth clustered storage architecture. It runs MapReduce for distributed data processing and works with structured and unstructured data (Figure 3).

**Figure 3:**
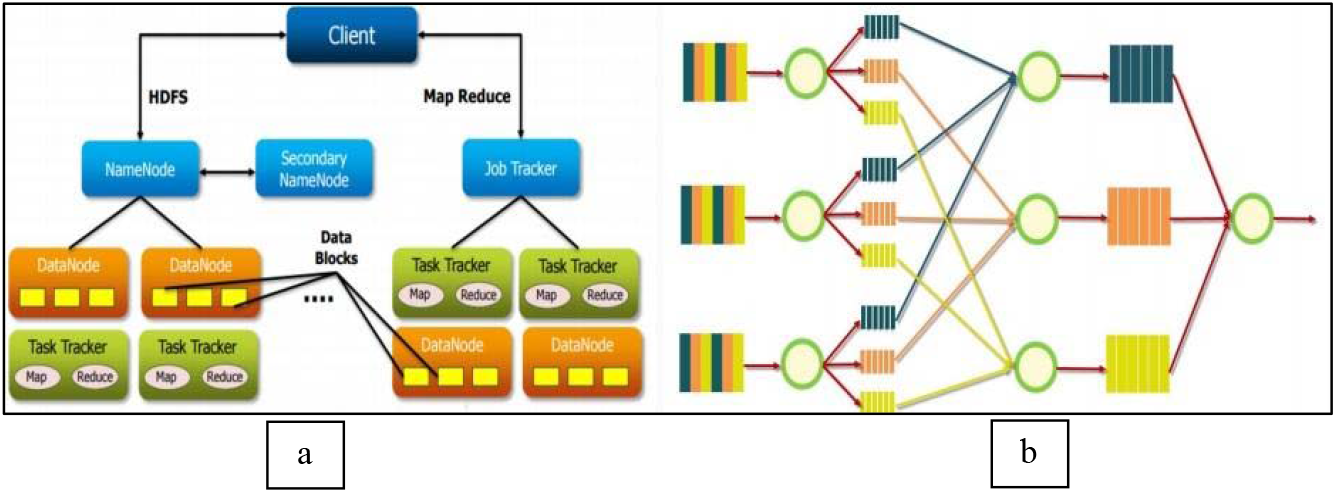
a) VIVIDHA: Hadoop Architecture b) MapReduce logic

#### 2.2.1 HDFS

HDFS is designed to reliably store very large files across machines in a large cluster. It stores each file as a sequence of blocks; all blocks in a file except the last block are of the same size. The blocks of a file are replicated for fault tolerance. Files in HDFS are write-once and have strictly one writer at any time. The NameNode makes all decisions regarding replication of blocks. It periodically receives a Heartbeat and a Blockreport from each of the DataNodes in the cluster. Receipt of a Heartbeat implies that the DataNode is functioning properly. A Blockreport contains a list of all blocks on a DataNode.

#### 2.2.2 NameNode

NameNode is the master node in the Apache Hadoop HDFS Architecture that maintains and manages the blocks present on the DataNodes (slave nodes). NameNode is a very highly available server that manages the File System Namespace and controls access to files by clients. HDFS architecture is built in such a way that the user data resides on DataNodes and never on the NameNode.

#### 2.2.3 DataNode

DataNodes are the slave nodes in HDFS. Unlike NameNode, DataNode is commodity hardware, that is, a non-expensive system which is not of high quality or high-availability. The DataNode is a block server that stores the data in the local file.

### 2.3 Web interface and variant file visualization

The web interface is based on the Java Server Faces (JSF) with Primeface and Javascript/jQuery (Figure 4). Server-side, the system is served by the JBoss Wildfly server on Ubuntu. Data is stored on the server side in user login.

**Figure 4:**
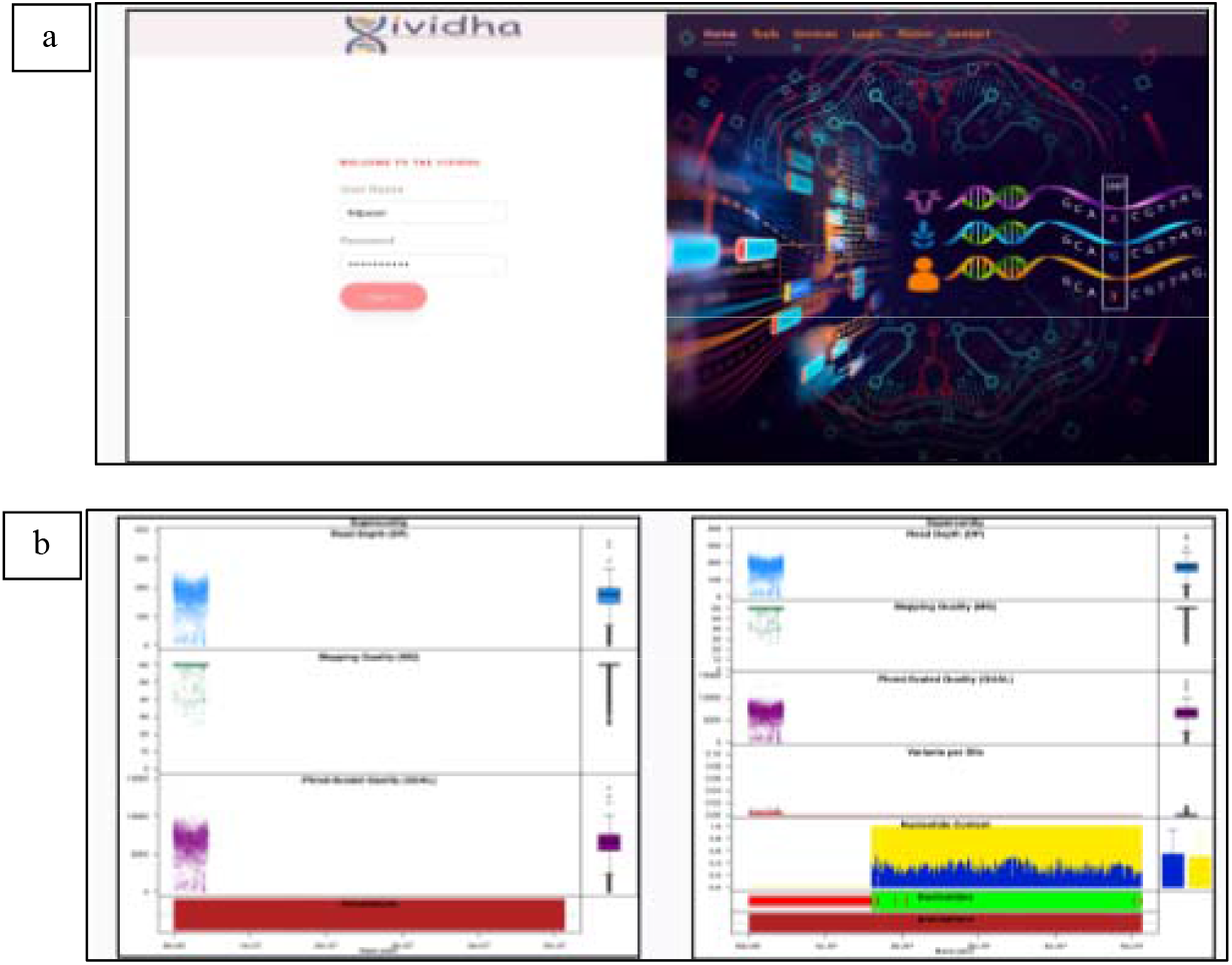
a) Web interface of VIVIDHA b) vcf file visualization

#### 2.3.1 Input Format

VIVIDHA accepts BAM file as Input. Files can be uploaded through SSH to the Hadoop cluster. User login is also created as a Hadoop client on Hadoop server. Prior to the submission of an analysis, a reference genome, tool name and suitable parameters can be selected from the drop down menu

#### 2.3.2 Visualization

VCF files are visualized using vcfR package [Knaus *et al*. 2017]. vcfR was designed to work on an individual chromosome, supercontig or contig.

#### 2.3.3 R Shiny

For web-based visualization R Shiny is used [URL: https://shiny.rstudio.com/]. Shiny is an R package that makes it easy to build interactive web apps from R.

## 3. Results and Discussion

With the increasing size of datasets, there is a dire need to develop scalable and cost-effective methodologies for variant calling. VIVIDHA is based on distribution of mapped files (BAM/SAM) into several overlapping chunks to enable fast computation without compromising on the quality of the called variants. Several callers have been employed on the sample dataset and a consensus strategy is used to arrive at the ‘optimal set of variants’. VIVIDHA enables variant discovery in low coverage data and reduce the overall ‘False Discovery Rate’. VIVIDHA has been developed using map-reduce on Hadoop wherein data is stored in a distributed manner using HDFS. The consensus methodology developed for vcf generation is based on the read length and mapping quality value.

### 3.1 Comparison with other tools

Comparison of various features in VIVIDHA with similar existing solutions like Halvade [Decap *et al*. 2015], Churchill [Kelly *et al*. 2015] and ADS-HCSPark [Xiao *et al*. 2019] was carried out (Table 1). Parabricks is a proprietary tool that is GPU-based and uses similar overlapping chunks of the alignment file for speed-up of variant calling (URL: https://resources.nvidia.com/c/tgen-success-story?x=sFVHf4&lx=OhKlSJ ; last accessed June 8, 2022). The comparison revealed that two functionalities viz., consensus variant calling and user interface for job submission are unique to VIVIDHA.

**Table 1:**
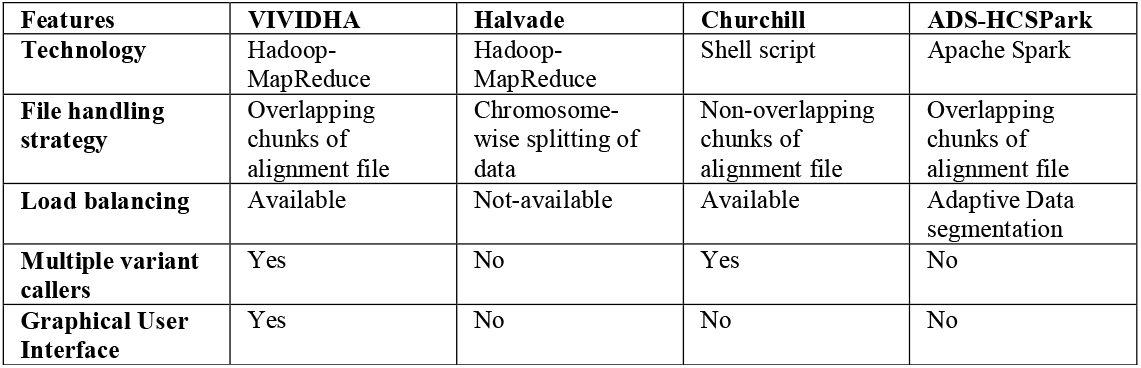
Comparison of VIVIDHA with four representative existing solutions.

| Features | VIVIDHA | Halvade | Churchill | ADS-HCSPark |
| --- | --- | --- | --- | --- |
| **Technology** | Hadoop-MapReduce | Hadoop-MapReduce | Shell script | Apache Spark |
| **File handling strategy** | Overlapping chunks of alignment file | Chromosome-wise splitting of data | Non-overlapping chunks of alignment file | Overlapping chunks of alignment file |
| **Load balancing** | Available | Not-available | Available | Adaptive Data segmentation |
| **Multiple variant callers** | Yes | No | Yes | No |
| **Graphical User Interface** | Yes | No | No | No |

*Salient features of VIVIDHA* (Figure 5)

- Use of overlap for splitting of alignment data
- Choice of variant callers as well as consensus calling
- User interface for ease of job submission
- Visualization of output using R

**Figure 5:**
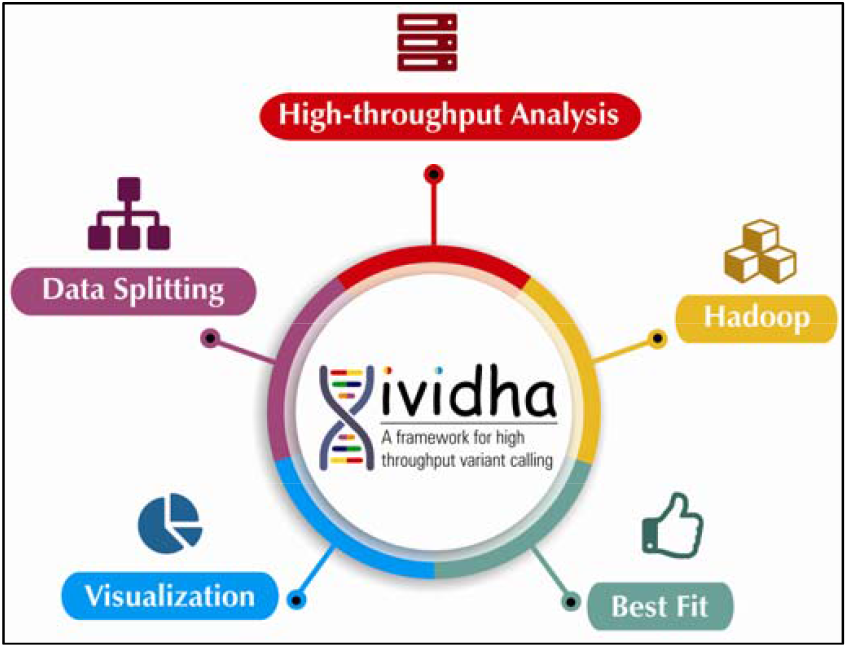
Features of VIVIDHA

### 3.2 Benchmarking

VIVIDHA has been benchmarked using three datasets from 1000 genomes project of varying sizes using GRCH38 as reference genome (Table 2). These datasets were benchmarked in triplicate on Ubuntu 18.04.5 LTS server with AMD EPYC 7452 32-Core Processor with 128GB RAM. Speed-up starting for 6x upto 10x was observed (Figure 6).

**Table 2:** Details of datasets used for benchmarking.

| Dataset | Size | Execution time for serial code (in minutes) | Execution time for Hadoop code (in minutes) | # SNPs |
| --- | --- | --- | --- | --- |
| NA18856 | 5.5GB | 436 | 73 | 1080790 |
| NA19116 | 18GB | 832 | 104 | 2386424 |
| NA19213 | 50GB | 1614 | 167 | 4654070 |

**Figure 6:**
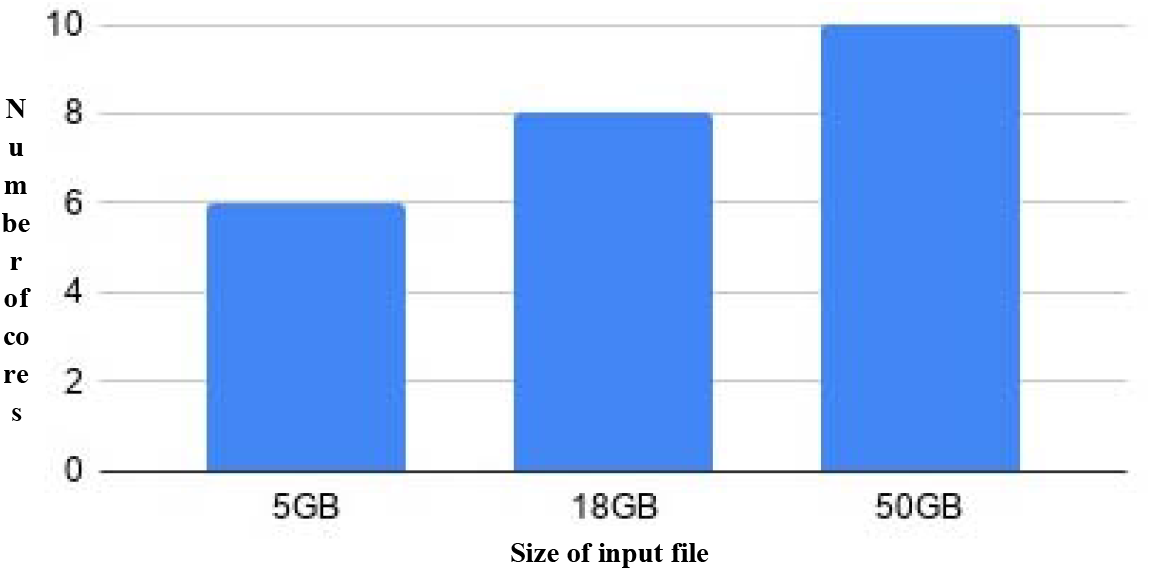
Plot of speed-up observed in VIVIDHA for different file sizes

### 3.3 Conclusion

VIVIDHA hides the complexities associated with variant prediction using different approaches by providing a very user-friendly interface for job submission as well as analysis. The consensus approach ensures prediction of actionable ‘high confidence variants’. The speed-up achieved using Hadoop-based overlapping window strategy of aligned genome file helps in timely analysis of clinical samples.

### 3.4 Availability

VIVIDHA is available as a stand-alone installer at URL: https://github.com/bioinformatics-cdac/VividhaInstaller

## Author Contributions

Software development: TB, RG and AS; Formal analysis: NB, SMK; Methodology: SMK and NB; Conceptualization: SMK and RJ; Data curation: NB; Writing - original draft: SMK, NB, TJ and AS; Writing - review & editing: RJ; Validation: NB, RG, AB and SJ.

## Acknowledgements

Authors acknowledge the Bioinformatics Resources and Applications Facility (BRAF) for the computing infrastructure. All the authors declare that they have no competing interests.

